# Genome-wide active enhancer identification using cell type-specific signatures of epigenomic activity

**DOI:** 10.1101/421230

**Authors:** Shalu Jhanwar, Stephan Ossowski, Jose Davila-Velderrain

**Affiliations:** Centre for Genomic Regulation (CRG), The Barcelona Institute of Science and Technology, Universitat Pompeu Fabra (UPF), Dr. Aiguader 88, Barcelona 08003, Spain; Developmental Genetics, Department of Biomedicine, University of Basel, Basel, Switzerland; Swiss Institute of Bioinformatics, Basel, Switzerland; Institute of Medical Genetics and Applied Genomics, University of Tübingen, Tübingen, Germany; Centro de Ciencias de la Complejidad (C3), Universidad Nacional Autónoma de México, Cd. Universitaria, México, D.F. 04510, México; Computer Science and Artificial Intelligence Laboratory, Massachusetts Institute of Technology, Cambridge, MA, USA; Broad Institute of MIT and Harvard, Cambridge, MA, USA

**Keywords:** Enhancer, cis-regulatory elements, regulatory genomics, epigenomic marks

## Abstract

Recently enhancers have emerged as key players regulating crucial mechanisms such as cell fate determination and establishment of spatiotemporal patterns of gene expression during development. Due to their functional and structural complexity, an accurate *in silico* identification of active enhancers under specific conditions remain challenging. We present a novel machine learning based method that derives epigenomic patterns exclusively from experimentally characterized active enhancers contrasted with a weighted set of non-enhancer genomic regions. We demonstrate better predictive performance over previous methods, as well as wide generalizability by identifying and annotating active enhancers genome-wide across different tissues/cell types in human and mouse.

## Introduction

Identification of active enhancers and their dynamics across different cell types, tissues and treatment specific conditions has become a prolific research area [1] [2]. Facilitated by the broad availability of high throughput sequencing techniques, the strategy for studying the molecular mechanisms underlying cell-fate determination has largely shifted from describing the expression profiles of lineage-specific transcription factors to characterizing the dynamic activity patterns of enhancers [3-5]. The context-dependent activity of enhancer sets and their disruption by mutation to a great extent underlie gene expression dynamics in development, cell differentiation, and disease progression [6] [7]. Enhancers are distal regulatory elements able to shape the cellular transcriptome by promoting the transcription of target genes [5]. Due to their functional relevance in normal development and diseases, the genome-wide identification of active enhancers from condition specific epigenomic data has emerged as a main goal and challenge in computational genomics [8]. Unlike other cis-regulatory elements (e.g., promoters), enhancers manifest heterogeneous structural and functional properties that are not completely understood [9–13]. Enhancers have specific features such as regulation of expression of distal genes irrespective of location or direction [2], low evolutionary conservation, non-coding transcription in the form of enhancer RNA [14] [15], variable motif composition, and functional dependence on chromatin conformation (e.g. enhancer-promoter interaction) specific to a cell or tissue [5], pose serious challenges for computational methods to accurately identify enhancers.

Previous studies have proposed computational methods that exploit genomic and epigenomic features to predict cis-regulatory elements [16]. To that end, the most commonly followed approach has been to assume an operational definition of enhancers based on (epi)genomic features known to be associated with them; for instance, genomic regions possess chromatin accessibility in addition to other regulatory features (e.g. DNaseI and P300 [17]), combinatorial epigenomic patterns of histone marks [18] [19], binding patterns of transcription factors (TFs) [20], or combinations of such features [21]. However, choosing one operational definition above others is not straightforward, and finding an optimal one based on associated features alone strongly depends on the state of knowledge and data availability at that time, both unavoidable limitations. Here, we propose an alternative unbiased approach to overcome such limitations.

Recent studies have generated unprecedented datasets of experimentally characterized enhancers using different functional approaches exploiting novel techniques. Kheradpour et al. have performed a massively parallel reporter assay to test the regulatory activity of ~ 2000 putative enhancers from two human cell-lines studied by ENCODE (K562 and HepG2) [22]. The FANTOM consortium recently provided a substantial resource of in vivo transcribed enhancers identified by sequencing of enhancer RNAs (eRNAs) using Cap Analysis of Gene Expression (CAGE) across a multitude of tissues and cell types in human and mouse [8] [14]. More recently, Kwasnieski et al. have experimentally tested enhancer activity of ENCODE predictions using CRE-seq in human K562 and H1 cell types [23]. Additionally, the VISTA database contains more than 2000 active enhancers characterized previously using reporter assays in human and mouse tissues [24]. The availability of diverse experimentally characterized enhancers provides an alternative to the use of imprecise and likely biased operational definitions of active enhancers based exclusively on correlated omics signatures.

Here, we present a machine learning (ML) based computational method for genome-wide prediction of cell- and tissue-specific active enhancers. The rationale behind the study is to learn exclusively from experimentally characterized active enhancers contrasted against a weighted set of non-enhancer background genomic regions, those epigenomic patterns with discriminatory power. We build, test, and apply the proposed method emphasizing four open problems in regulatory (epi)genomics: whether (1) robust signatures of enhancer activity can be learned from an integrative set of enhancers whose activity has been experimentally characterized using different techniques (2) uncovered patterns can be used to accurately predict enhancer activity genome-wide (3) the predictive power is independent of the cellular conditions where the discriminatory signatures were first derived from and (4) the trained ML model generalizes to different tissues and species. We explore such problems following a supervised ML approach and provide a publicly available tool termed as **G**eneralized **E**nhancer **P**redictor (GEP). The computational framework is available at https://github.com/ShaluJhanwar/GEP.

## Results

### An integrative training dataset

In order to extract patterns of epigenomic features discriminating active enhancers, we have assembled a training dataset of experimentally characterized active enhancers corresponding to HepG2 and K562 human cell types (*n* = 2, 128) from Kheradpour et al. [22] (Table S1.A, additional file) and a balanced set (*n* = 2, 128) of genomic elements with no enhancer activity including promoter, gene-body, and heterochromatin regions with an approximate 5:3:2 ratio, respectively (see Methods). We have characterized instances of the training set (Table S1.B, Additional file) with 16 features describing the epigenomic state of the corresponding genomic region. Considering that enhancer and promoter elements share similar characteristic properties [14] [25], we have decided to include a higher relative number of promoters in the negative class instead of the commonly used random background sampling strategy to reduce the chance of prediction errors, i.e. prediction of promoters as enhancers.

### Feature selection and model evaluation

We have chosen Random Forest (RF) and Support Vector Machine (SVM) classifiers to build predictive models for classification of active enhancers. In order to select informative features for accurate prediction, we estimated the relative importance of the individual features using the RF algorithm (see Methods). Figure 1B shows the resulting feature ranking. As expected, the top-ranked epigenomic features correspond to chromatin marks (H3K4me1 and H3K27ac) commonly associated with enhancer activity [26] [27]. H3K4me3 is known to be more frequent on promoters than enhancers, and H3K4me1 is more frequent on Enhancers; hence we explored these properties as ratio of histones. Interestingly, these non-conventional features such as the ratio of histones (H3K4me1/H3K4me3) and distance to the nearest annotated transcription start site (TSS) are highly discriminatory as well (top 4 rank). Based on the feature ranking in several rounds of training and testing, as well as the broad availability of the histone marks in large studies such as ENCODE and NIH Roadmap Epigenomics [28], we selected a minimal set of nine features for model training that includes the top seven ranked features (Figure 1B) in addition to the two frequently interrogated marks associated with gene transcription (H3K36me3) and repression (H3K27me3). The model with these nine features was found to be as informative as the model with all 16 features (see below).

**Figure 1:**
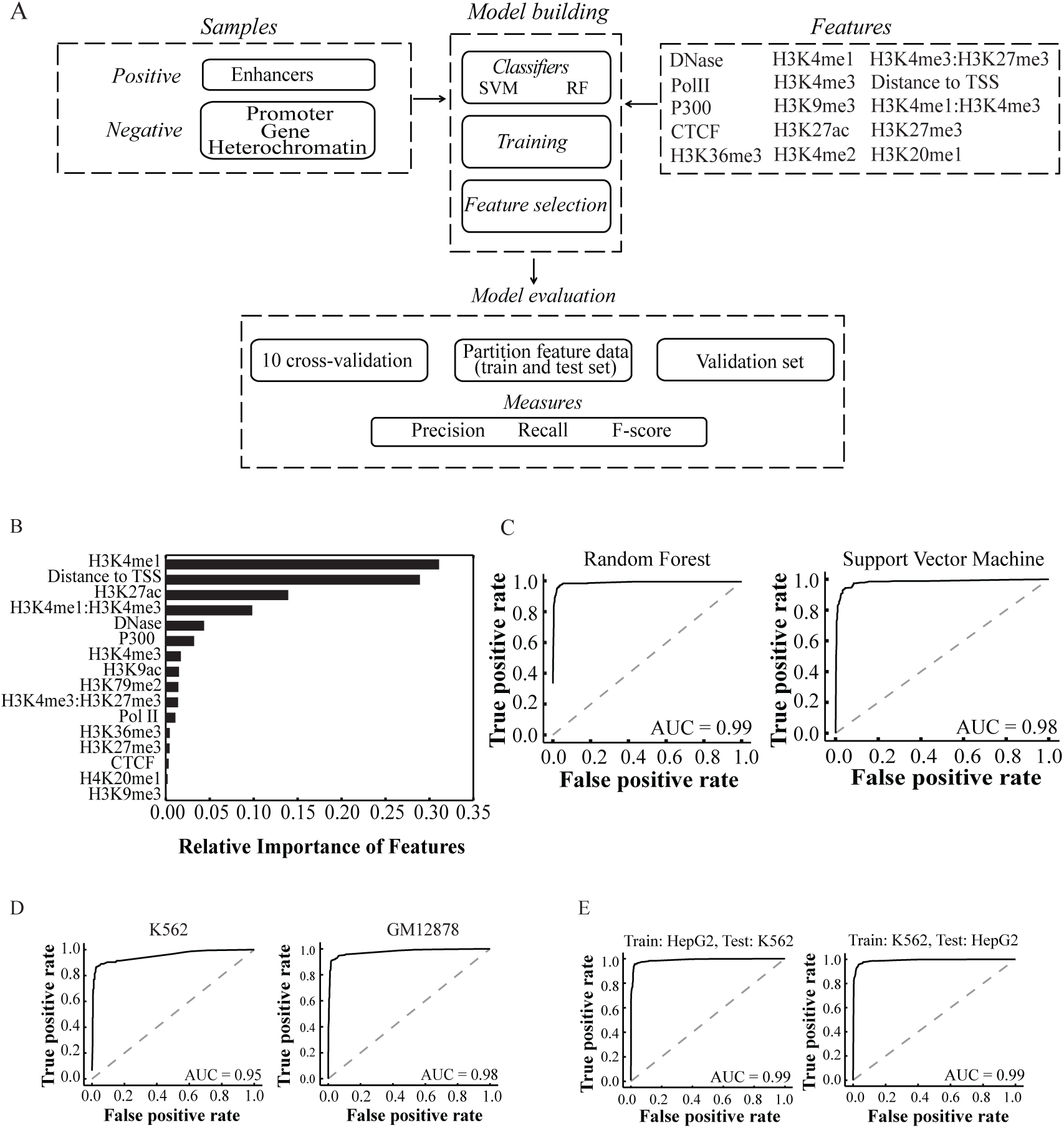
Workflow and validation of GEP. A) Machine-learning framework used for training, feature selection and evaluation of GEP B) Relative importance of (epi)genomic features using Random Forest. Black bars represent the relative importance of the features for distinguishing enhancer from non-enhancer class during training of the GEP model from experimentally validated enhancers in K562 and HepG2 C) ROC curve of the classifiers when train on 80% of the training data and test on 20% of the training data D) ROC curve on validation of FANTOM5 in vivo transcribed enhancer sets using random forest E) ROC curves with random forest in order to test independence of cell types during enhancer prediction.

For evaluation of the model trained on HepG2 and K562 human cell types, we performed three experiments. First, in 10-fold stratified cross-validation the mean test score was 0.98, showing little variability across the 10-folds (Table S2A, Figure S1A). Second, we split the training dataset into a train (80%) and a test (20%) set. Both the classifiers (RF and SVM) have achieved > 93% of accuracy on the test set (Table S2B, Figure 1C). A convergence of the predictive accuracy in the learning curve as shown in Figure S1C suggested that the amount of training data is sufficient for learning discriminatory combinations of the nine features. In the third and final experiment, to further evaluate the predictive performance of the model, we used two independent validation datasets of the in vivo transcribed enhancers corresponding to K562 and GM12878 cell-lines (FANTOM5) expressed in at least two replicates. Interestingly, the models have showed high performance, with the RF performing better (F-score > 0.90) than the SVM (F-score > 0.80) (Table S2C; Table S3; Figure 1D and Figure S1.B). Recent studies have highlighted the relevance of sequence-based features in enhancer identification [16]. We tested whether including additional genomic features such as the presence of CpG Islands, evolutionary (PhastCons) and TFBS conservation score (ENCODE), GC content, and TF binding motifs (Jaspar and PreDREM) might improve the performance of GEP using a gold-standard benchmarking dataset of experimentally characterized enhancers (see Benchmarking section). The resulting model have showed a slightly lower performance (0.54 accuracy and 0.54 F-score) with respect to the original GEP model (0.56 accuracy and 0.55 F-score (see Benchmarking section; Table S6), suggesting that sequence-based and evolutionary signatures do not provide additional discriminatory information using the proposed machine-learning model. Based on the above results and the substantially shorter training time of RF compared to SVM, the final GEP model consists of a RF trained on eight epigenomic features and one genome annotation feature (distance to annotated TSS) corresponding to the sequences with enhancer activity, previously characterized in HepG2 and K562 cell types. The design of the ML framework (GEP) is illustrated in Figure 1A.

### Model performance is independent of cell type

In order to test the dependence of the method on the choice and specifics of the cell types used for training, we performed two experiments, 1) we trained the model on K562 data only and tested it on HepG2 and 2) vice-a-versa. In both the experiments, the trained model have showed excellent performance on the test sets (*∼*96% accuracy and 0.96 F-score) (Figure 1E). Moreover, we tested GEP performance on a set of enhancers known to be active (in vivo transcribed, FANTOM5) in other cell type that was not used for training *i.e* GM12878. GEP showed a remarkable performance of 98% AUC on GM12878 (see Table S2C; S3 and Figure 1C). These results indicate that GEP performs in a cell type independent manner. The genome-wide prediction of enhancers across different cell types/tissue in mammals reported and annotated herein further strengthened this conclusion (see below).

### Application of GEP for genome-wide enhancer identification in mammals

In order to accurately and efficiently identify active enhancers genome-wide, we have implemented the following protocol: (1) the entire genome is fragmented into 500bp windows using a step size of 250bp (2) a subset of “active” genomic regions is pre-selected based on acetylation marks [29][30] (3) a set of annotated non-enhancer cis-regulatory elements (i.e. promoters annotated in GENCODE TSS) is removed from downstream analyses (4) each of the remaining active regions is classified into “enhancer” or “non-enhancer” using GEP (5) putative enhancer regions are ‘shrunk’ based on three enhancer associated epigenomic marks (*i.e.* the upstream and downstream borders are shifted until a position with overlap of at least one of DNaseI, H3K4me1 and H3K27ac is reached) and (6) enhancer predictions with overlapping boundaries are clustered in order to define the final boundaries of putative enhancers (Figure 2A). To functionally annotate these *insilico* predicted enhancers, we have defined an activity score based on the overlap of regulatory activity signals (see Methods). Briefly, we considered the overlap of predicted enhancers with five individual categories of independent data as evidence suggestive of regulatory activity; the data includes chromatin accessibility (DNaseI), TF binding potential as TF-ChIP-seq, TFBS motifs, experimentally validated enhancers (VISTA and FANTOM5) and long-range chromatin interactions (Hi-C and ChIA-PET).

**Figure 2:**
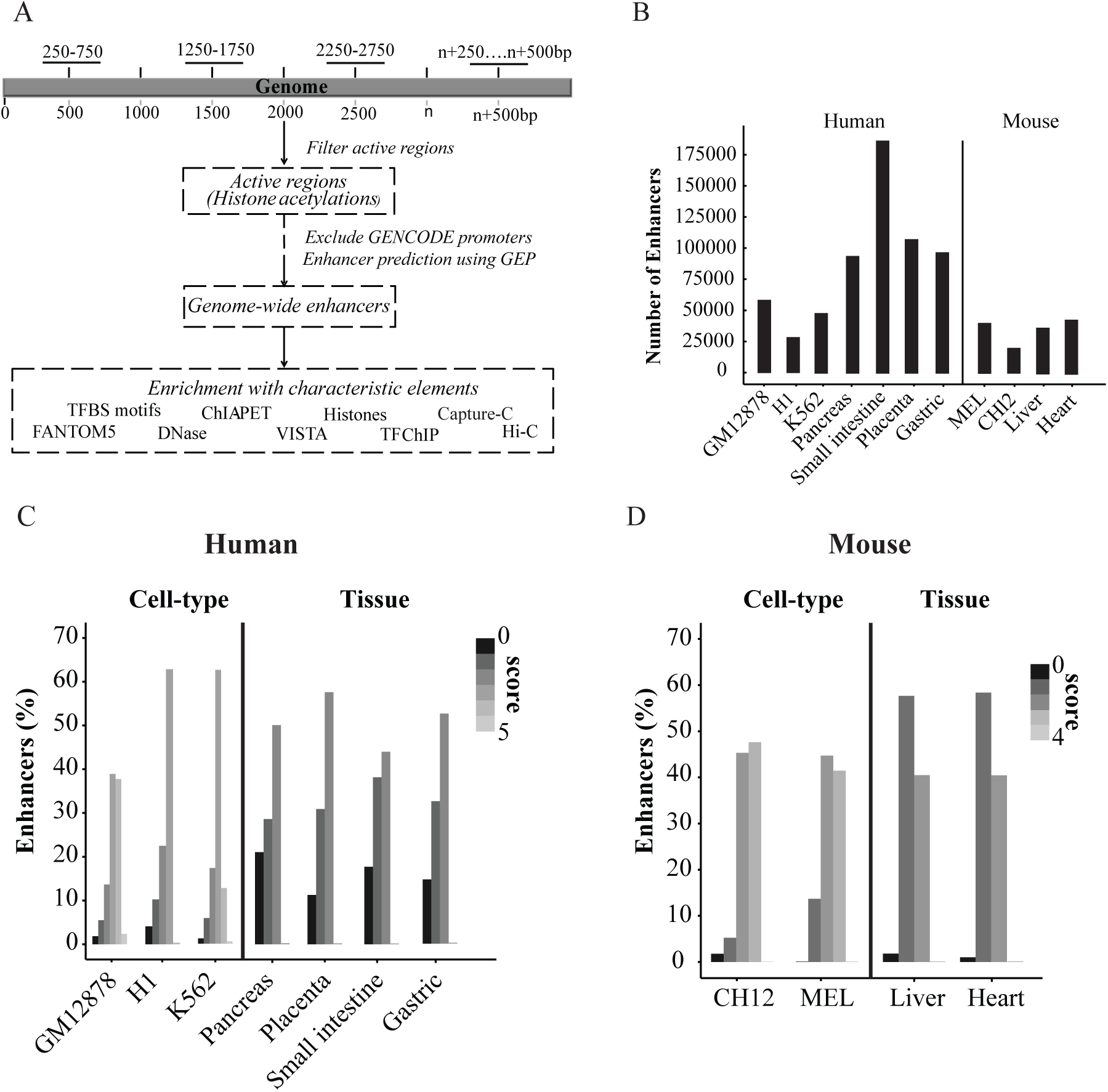
Genome-wide identification of enhancers in mammals using GEP *Insilico* genome-wide identification and evaluation of active enhancers using GEP B) Number of *insilico* predicted putative enhancers across different cell-/tissue types in human and mouse. Activity scoring of predicted enhancers in C) human cell types and tissues and D) mouse cell types and tissues. The overlap of active elements with suggestive of enhancer activity that includes DNaseI, active histone marks, TF-ChIP, TFBS motifs, chromatin interaction (ultra-deep HiC and ChIA-PET) and experimental evidences of enhancers from different resources.

Following the steps described above, we first performed genome-wide enhancer prediction and activity scoring in three commonly used human cell-lines. A total of 48,598 (*∼*1.29% of genome with 594bp median size), 58,857 (1.81% of the genome with 599bp median size), and 28,821 (0.36% of the genome and 274bp median size) putative enhancers were found to be active in K562, GM12878 and H1 respectively. Activity scoring have revealed that the majority of the enhancers (> 95%) were supported by at least one independent category of regulatory activity across all the cell types, > 63% being supported by at least three (Figure 2C, Table S5). Further, we have compared the predictions with the regulatory region categories proposed by Yip et al. [20] based on the binding events (ChIP-seq) corresponding to more than 100 TFs. Here, more than 75% of GEP’s enhancer predictions fell either into the category of ‘binding active region’ (BAR) or ‘distal regulatory module’ (DRM) across all the cell types (Figure S2A). Only a small fraction (*<* 1 %) overlapped with the ‘proximal regulatory module’ (PRM), suggesting that most GEP predictions are not promoters. As expected, the predicted enhancers were found to be far from TSS with median distances ranging from 5123.5bp to 9708bp depending on the cell type (Figure S2B).

In order to predict enhancers in primary human tissues, we have exploited the large epigemonic dataset released by the NIH Roadmap Epigenomics consortium [28]. Given that P300 is not interrogated in most of the tissues, we trained a new RF model using the same training data and features as described above, but omitting P300 as a feature. The removal of P300 has negligibly affected the performance of the model as compared to the GEP model trained with P300 (Table S2A). Hence, the new model relies only on the histone marks included in the ‘core epigenome’ defined by Roadmap Epigenomics that are also present in the majority of ENCODE cell types [31], mouseEncode cell-lines and tissues [32], and BLUEPRINT cell types [33], making the model broadly applicable. Using the new model without P300, we performed genome-wide predictions of enhancers active in placenta, pancreas, gastric and small intestine human tissues. A total of 1.05% (pancreas; 94,453 enhancers), 1.17% (gastric; 97,504 enhancers), 2.38% (small intestine; 187,097 enhancers) and 2.62% (placenta; 107,974 enhancers) of the human genome were predicted as active (Figure 2B) enhancers. For activity scoring of enhancers predicted with Roadmap data, limited data availability has precluded the consideration of chromatin interaction and TF binding events. However, we included tissue-specific data corresponding to in vivo transcribed enhancers from FANTOM5 for additional support. Notwithstanding data limitations, a majority of predicted enhancers (> 80%) active in each of the tissues were supported by at least one regulatory signal (Figure 2C; Table S5).

Recent studies have shown a high conservation of epigenomic signals between regulatory elements in human and mouse [34]. Following the assumption, we have used GEP trained exclusively on human data to predict active enhancers in mouse. We performed genome-wide enhancer identification on the mouse cell types MEL (analogous to K562) and CH12.LX (analogous to GM12878), and tissues (heart and liver). We predicted 1.70% (CH12; 20,747 enhancers), 2.09% (MEL; 40,750 enhancers), 2.74% (Heart; 44,153 enhancers) and 2.40% (Liver; 37,466 enhancers) of the genome (Figure 2D; Table S8) possessing enhancer activity. Although cell-/tissue-type-specific data is less abundant for mouse than human, still a majority of the predictions (> 95%) were supported by more than one independent signal associated with regulatory activity including open chromatin, TF-binding, and TFBS motif features (Figure 2C and Figure 2D; Table S5). Overall, the results indicate that the predictive performance of the GEP model trained on two human cell-lines is able to generalize to independent human cell types and tissues, as well as to other mammalian model systems (mouse).

### Benchmarking GEP against existing enhancer prediction methods

Previously reported and widely accepted ML based enhancer predictors include CSI-ANN [32], RFECS [12], DEEP [16], ChromHMM [34] and Segway [14]. We have performed benchmarking with pre-existing methods at two broad levels using empirical evidence of regulatory activity as evaluation categories: a) a general signatures of chromatin and regulatory activity that included epigenomic marks suggestive of regulatory activity (P300, DNaseI and TF-ChIP), expressed enhancers (eRNA) in K562 (FANTOM5), evidence with recently available 3D regulatory interactions (Hi-C and ChIA-PET) and False Positive Rate (FPR) detection using CAGE promoters specific to K562 (FANTOM5) b) specific signatures of enhancer activity using a comprehensive gold-standard set of experimentally validated enhancers from different resources. The dataset included 1,621 non-active enhancer elements (CRE-seq assay) and 2,392 active enhancers including VISTA [20] (reporter assay), FANTOM5 [18] (CAGE) and Kwasnieski et al., [37] (CRE-seq with P-value *<* 0.05).

When benchmarked using general signatures of regulatory potential, GEP has showed the best performance among all, except one (second place). On the basis of regulatory activity based on histone acetylations, TF binding ChIP, open-chromatin and P300 binding (Table 1), GEP showed the best performance (F-score 34.07), followed by Segway (F-score 33.26). In regulatory 3D-interactions analysis based on ChIA-PET and Hi-C data (Table 1), RFECS showed the best PPV (17.8) followed by GEP (17.4), supporting the good quality of predictions. In the case of promoter overlap (i.e., false positive rate in CAGE promoter overlap), ChromHMM showed least overlap with K562 specific TSS from FANTOM5. CSI-ANN and DEEP showed a greater overlapped with CAGE promoters (Table 2) relative to other methods, suggesting their predictions were not enhancer specific. Furthermore, with K562 in vivo transcribed enhancer (FANTOM5) expressed in at least 2 replicates (Table 4), GEP has over performed other methods (75.70% base-pair overlap and 83.94 numbers overlap). Interestingly, when benchmarked with the specific signatures of regulatory potential (gold-standard experimentally characterized enhancers), GEP outperformed other methods (F-score 0.55) followed by DEEP (F-score 0.53) as shown in Table 3. Note that the data included in the benchmarking set were not used during the training phase of the model.

**Table 1:**
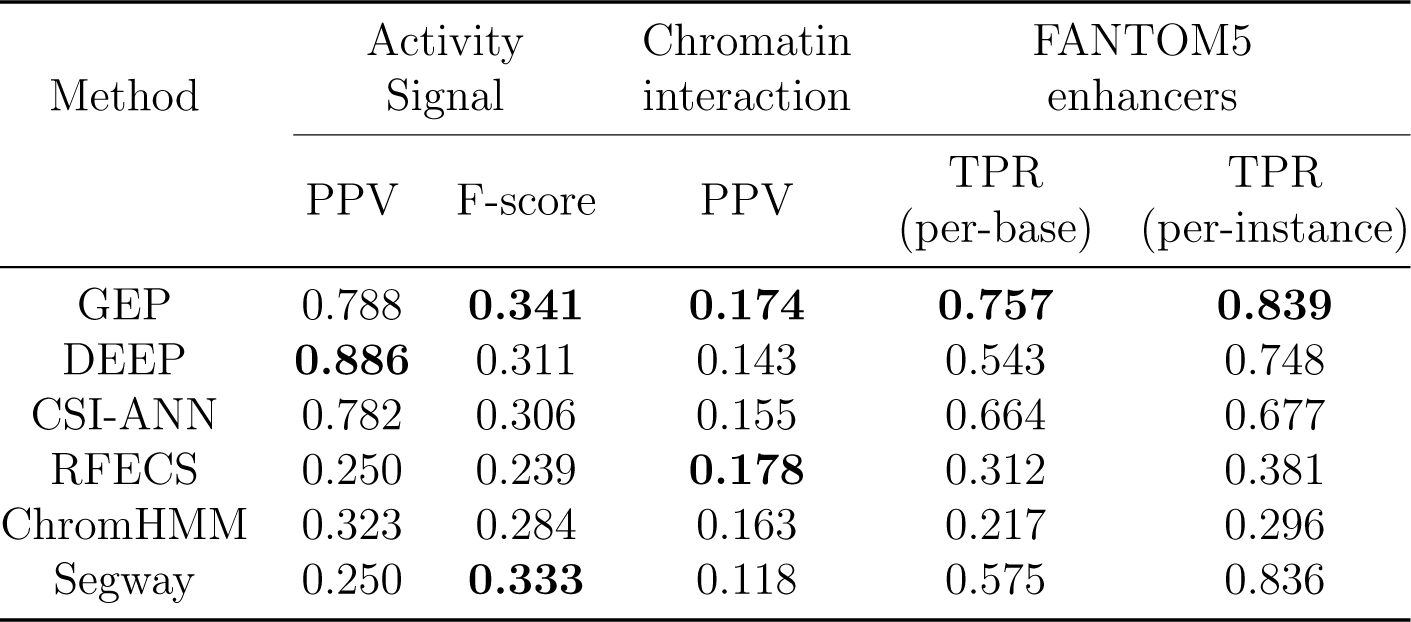
Benchmarking of GEP with existing ML tools. Activity signal is based on the overlap with the union of regulatory activity signatures in K562, including DNaseI, TF-ChIP, P300 and histone acetylation. PPV and F-score have been calculated per base, with positions in predicted enhancers covered by at least one activity feature considered as true positive, else as false positives. Chromatin interaction is based on the overlap of predicted enhancer with chromatin interaction loops obtained via ultra-deep Hi-C or ChIA-PET in K562 (minimum of 1bp overlap required per instance). FANTOM enhancers TPR (sensitivity) is based on the recall of expressed K562 enhancers (significant CAGE signal in at least 2 replicates) annotated by the FANTOM5 consortium

| Method | Activity Signal |  | Chromatin interaction | FANTOM5 enhancers |  |
| --- | --- | --- | --- | --- | --- |
|  | PPV | F-score | PPV | TPR (per-base) | TPR (per-instance) |
| GEP | 0.788 | **0.341** | **0.174** | **0.757** | **0.839** |
| DEEP | **0.886** | 0.311 | 0.143 | 0.543 | 0.748 |
| CSI-ANN | 0.782 | 0.306 | 0.155 | 0.664 | 0.677 |
| RFECS | 0.250 | 0.239 | **0.178** | 0.312 | 0.381 |
| ChromHMM | 0.323 | 0.284 | 0.163 | 0.217 | 0.296 |
| Segway | 0.250 | **0.333** | 0.118 | 0.575 | 0.836 |

**Table 2:**
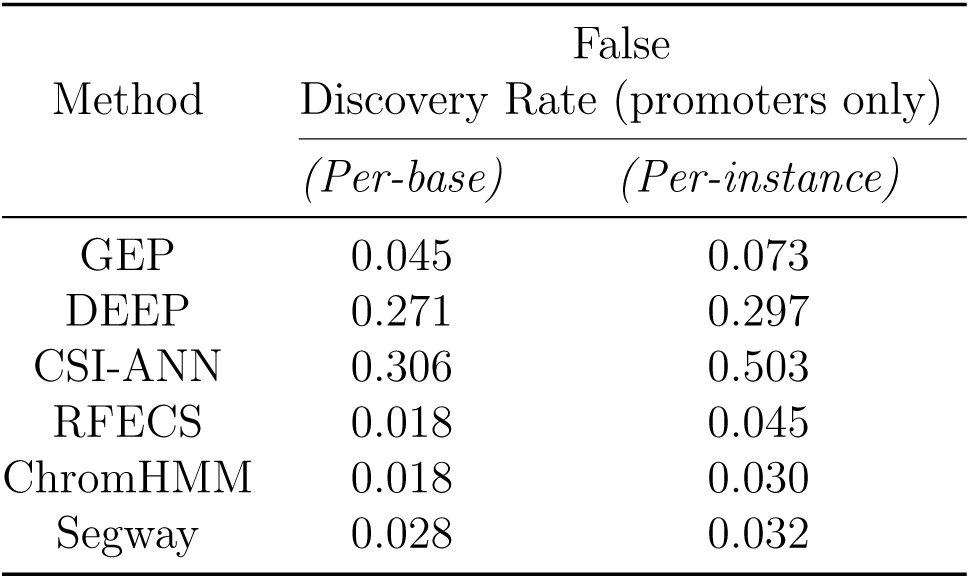
Benchmarking of GEP based on promoter overlap. The overlap of predicted enhancers with regions identified as active promoters defined by CAGE-seq of K562 (−1000 bp to +500bp to TSS), was used as a false positive definition.

**Table 3:** Benchmarking of GEP based on experimentally characterized active and inactive enhancers. The dataset of experimentally characterized enhancers were obtained from different studies and resources (Kwasnieski et al. [23], VISTA, and FANTOM5). The number as well as percentage of genome identified as enhancers varied across different methods.

| Method | Approach | Number<br>of<br>enhancers | Percentage of hg19<br>covered (without chrY) | F-score |
| --- | --- | --- | --- | --- |
| GEP | Supervised | 48,598 | 0.0013 | 0.55 |
| DEEP | Supervised | 151,500 | 0.0100 | 0.53 |
| CSI-ANN | Supervised | 17,309 | 0.0114 | 0.5 |
| RFECS | Supervised | 65,329 | 0.0430 | 0.46 |
| ChromHMM | Semi-supervised | 205,137 | 0.0367 | 0.43 |
| Segway | Unsupervised | 1,237,490 | 0.0931 | 0.5 |

Taken together, the benchmarking results (Table 1-3) demonstrated that GEP either outperformed or showed similar performance as other tools at both computationally as well as experimentally characterized level, implying the importance of GEP as a novel and accurate enhancer predictor.

## Discussion

Previous studies have demonstrated the use of combinatorial patterns of epigenomic signals to determine distal genomic regions likely to have context specific regulatory activity [2]. More recently, a growing body of studies has started to experimentally characterize a large sets of enhancers active under certain conditions. By integrating such studies, we were able to assemble a comprehensive repertoire of active enhancers, useful for training, validation and benchmarking of *insilico* enhancer prediction methods. Instead of using approximate definitions of active enhancers based on single or a combination of chromatin marks, here we have derived the epigenomic patterns informative of enhancer activity exclusively from experimentally tested genomic elements with enhancer activity. Additionally, we have structured the genomic background in a balanced way representative of potential non-enhancer regulatory elements (promoters) and gene body regions, in addition to the randomly sampled genomic regions. We have designed a novel supervised Random Forest based model (GEP) trained on combinatorial epigenomic activity patterns derived from human cell types (HepG2 and K562).

The evaluation of the proposed workflow on the different test and validation datasets across different cell types reinforced the satisfactory performance of GEP. Furthermore, a comprehensive benchmarking of GEP against pre-existing enhancer prediction methods based on the consistency of predictions with other indicators of regulatory activity (DHS, P300, TFBS-ChIP and histone acetylations), promoter-enhancer interaction (ChIA-PET and ultra-deep Hi-C), enhancer expression (eRNA), and CAGE-based promoter prediction (FANTOM5) have demonstrated the improved performance of GEP over existing methods. Moreover, we have assembled a gold-standard set of thousands of experimentally active and non-active enhancers using diverse functional techniques (i.e., CAGE, CRE-seq, and reporter assay), which may prove useful for future studies approaching the (epi)genomic characterization of active enhancers, as well as their defining features, both open problems of broad interest [13].

Unfortunately, except for the most commonly studied Encode cell types, the availability of epigenomic data and validated enhancers for human cell types and tissues are scarce. Considering this issue, we built the GEP model in a way that it depends only on the commonly available features for the majority of the cell/tissues types included in international projects such as ENCODE [31], mouseENCODE [32], Roadmap Epigenomics [28] and Blueprint [33]. The working of GEP relied merely on a set of histone and chromatin marks defined as the ‘core epigenome’ by the Roadmap Epigenomics consortium [43].

Interestingly, during the building and testing phase of the model, we found that sequence-based features (e.g., motif enrichment and sequence conservation) did not improve the performance of GEP, with the exception of distance to annotated TSS. This observation may provide hints for future studies associated with the molecular mechanisms involved in the functional diversification of regulatory elements, a topic of growing interest [13] [35]. It is consistent with the recent results suggesting that conservation at the level of function (e.g. motif composition) might be a common mode of enhancer evolution, rather than conservation at a sequence level. [36].

The main goal of computational classifiers is to provide a model able to predict the class of previously unobserved instances. We demonstrated that GEP’s satisfactory predictive performance generalizes across human cell types independent of the training cell types. Furthermore, based on the growing evidence for highly conserved general mechanisms of epigenomic regulation, at least across mammals [37], we hypothesized that discriminatory patterns of enhancer activity derived from specific human cell lines can be utilized to predict active enhancers in any cell/tissue type across mammalian species. If proven correct, the utility of tools such as GEP will increase substantially, given that extensive experimental data for model training has been produced for a few human cell lines (GM12878, K562), but not for other human cell types and tissues in humans and other species. As a proof of principle, and in order to provide support to our claim, we performed genome-wide enhancer prediction in human and mouse cell types and tissues using a single GEP enhancer model trained on K562 and HepG2 cell lines. Importantly, to the best of our knowledge, we showed for the first time that the patterns computationally derived from one species can be used to make genome-wide predictions of active enhancers in another species. These results are consistent with the hypothesis of a conserved epigenomic code responsible for establishing gene expression patterns in a tissue and developmental stage specific manner, which also aligns with recent discoveries in what has been referred as” comparative epigenomics” [38]. We have explored multiple independent signals for enhancer activity (comprising genomic, epigenomic, transcription regulation and chromatin structure features) to support the regulatory activity of the majority of predictions in both human and mouse. Note that by definition, ML based methods are limited by the amount of data, information and knowledge available at the time of training. As such, future conceptual developments and new experimental datasets related to the mechanisms by which distant cis-regulatory elements work is likely to improve computational methods, including GEP. For example, some results suggest that the activity status of regulatory elements may be subject to graduation, or to belong to discrete classes of activity; for example primed activity [39] functionally defined by eRNA [40]. However, the data and knowledge available to date do not enable a clear-cut definition and representation of regulatory activity at that resolution, thus precluding the formulation of more specific predictive models. Likewise, further epigenoimc data from different mammalian species is needed in order to further support our hypothesis of epigenomic inter-species conservation and its utility for regulatory element prediction.

## Conclusions

We have built; tested and applied a novel ML based method to perform genome-wide identification of active enhancers. The publicly available method GEP has been trained on human cell-lines and can be readily applied for genome-wide prediction in mammalian species following the protocol in Figure 2A, given that the required ‘core’ histone marks and chromatin accessibility information (DNaseI or ATAC-seq) is available for the tissue or cell type of interest. Unlike previous methods, the GEP model has been trained exclusively from experimentally characterized enhancers and a heterogeneous balanced background dataset. Benchmarking of GEP with pre-existing computational methods, as well as genome-wide predictions and annotation in different conditions and species demonstrate the practical relevance of the method. A direct application of our method is the identification of genomic regions, which if mutated (regulatory alterations), would likely have a significant phenotypic effect that might be directly linked with the manifestation of complex diseases originating from specific tissues and organs.

## Methods

### Raw dataset

Peak calling data used in this study for human cell-lines (K562, HepG2, GM12878, H1) and mouse cell-/tissue types (MEL, CH12.LX) were retrieved from the ENCODE and mouse ENCODE download portal [44]. Processed peak calling data for enhancer identification in human tissues were downloaded from the NIH Roadmap Epigenomics Project [43]. In each case, the pre-processed data for 6 histone modifications (H3K4me1, H3K4me3, H3K27ac, H3K36me3, H3K9ac, H3K27me3), open-chromatin (DNaseI), and TFBS ChIP-seq corresponding to cell types and tissues were used in the analysis. All used resources are listed in Table S9 (additional file 1). The dataset used for testing of the applicability of the model in independent cell (tissue)-types in human and mouse was downloaded from the ENCODE and Roadmap Epigenomics data portals. The data sources corresponding to each cell (tissue)-type are provided in Table S9.

### Training dataset

The training dataset represents positive and negative classes in 1:1 ratio. Classes where chosen as follows:

- Positive class: active enhancers were taken from [22], where regulatory activity in predicted human enhancers from ENCODE cell-lines HepG2 and K562 were experimentally tested using massively parallel reporter assays. Each instance, originally of 145 bp (Table S1.A; Additional file 1), was transformed into corresponding regions of 500bp by considering equally sized flanking sites. The 500bp size was chosen based on the average size of enhancers active in K562 cell-line (at least 2 replicates) according to the FANTOM5 (Figure S3) enhancer atlas [14]. A total of 1,296 and 832 instances of active enhancers corresponding to HepG2 and K562 cell-lines, respectively, were taken from [22]. Corresponding genomic coordinates were given in Table S1.B (additional file 1).
- Negative class: Non-enhancer cis-regulatory elements (i.e., promoters), exons, introns and intergenic regions were taken in defined shares deviating from their genomic fractions as negative instances for training purposes. The heterogeneous set was selected following a random sampling procedure subjected to the following constraints: (1) genome-wide location of the instances follows the same distribution across chromosomes as the one presented by the enhancer set; and (2) the whole negative set is comprised of promoter, gene body and heterochromatin elements with approximate proportions of 50, 30 and 20 per cent; respectively. Positive and negative training classes were assembled independently for each cell-line (HepG2 and K562) and subsequently merged to obtain a single training dataset with a total of 2,128 positive (negative) instances (Table S1.B, additional file 1). All annotations were based on human Gencode v19 and mouse vM1 [45] corresponding to genome assembly versions hg19 and mm9 respectively.

### Feature Extraction and Selection

A set of features was selected in order to numerically characterize the state of each instance. The set contains varied types of features including: histone modifications, chromatin accessibility, DNA binding proteins, ratio of chromatin marks (H3K4me1*/*H3K4me3 and H3K4me3*/*H3K27me3), and the distance to the nearest TSS. For each feature based on ChIPseq data, the corresponding peak files were downloaded from ENCODE [44]. Details on data resources are listed in Table S9 (additional file 1). A feature ranking approach was followed as implemented within the random forest algorithm in scikit-learn v0.16.0 python library, setting the best parameter value of n estimators to 100.

### Model Building and Evaluation

The models were trained using two popular ML classifier algorithms i.e. Random Forest and Support Vector Machine, as implemented in the python library scikit-learn v. 0.16.0. Parameter optimization was performed based on the 10-fold cross validation performance. For model validation, in vivo transcribed enhancers in K562 and GM12878 cell-lines were taken from FANTOM5 [46] and used as independent test sets. Only enhancers expressed in at least two replicates were considered as active in a particular cell-line. Model performance was evaluated using standard ML-metrics i.e. accuracy, F-score, and Area under the ROC curve (AUC).

### Computational Validation of Genome-wide Predictions

In order to provide support for potential regulatory activity, genome-wide enhancer predictions were characterized based on (epi)genomic features known to be associated with active enhancers. We used a simple activity scoring method for predicted enhancers based on the evidences of overlapping with independent signals of regulatory activity. To this end, available enhancer associated elements were divided into maximum of five categories, based on the common characteristics. Each category of signals can be present or absent in a specific enhancer, and the sum of present signal categories represents the confidence score with a maximum of five. The proposed categories of enhancer activity associated elements were i) overlap with DNaseI, H3K4me1 or H3K4me3 ii) annotated TFBS motifs from JASPAR [47] and PreDREM [48] iii) experimental validation, including enhancer sets validated using CRE-Seq [23], CAGE [14] and reporter assay (VISTA [24]) iv) ChIP-seq of transcription factors [49] regulatory modules proposed by Yip et al. based on ChIP-seq of more than 100 TFs v) long-range chromatin interactions of promoters and enhancers (loops) including predictions based on ChIA-PET [41] and ultra-deep Hi-C [42]. Details of the datasets used for functional characterization purposes are provided in Table S9 (additional File 1).

### Benchmark with existing prediction methods

In order to compare GEP prediction accuracy with those of previously published methods including CSI-ANN [32], RFECS [12], DEEP [16], ChromHMM [34] and Segway [14], publicly available genome-wide enhancer predictions corresponding to individual methods for myelogenous leukemia K562 cell-line (hg19 genome) were downloaded. ChromHMM and Segway predictions were obtained from ENCODE [50], whereas genome-wide predictions of K562 given by DEEP, RFECS and CSI-ANN were obtained from DEEP [51]. Using empirical evidence of regulatory activity as evaluation categories, a bench-marking of GEP with existing methods has been performed at two broad levels:

a) A general signatures of chromatin and regulatory activity - this category includes epigenomic marks suggestive of regulatory activity, expressed enhancers (eRNA) in K562 (FANTOM5) and evidence with recently available 3D regulatory interactions (Hi-C and ChIA-PET). Additionally, CAGE promoters specific to K562 (FANTOM5) was considered to calculate False Positive Rate (FPR) of predictions. Due to the variations in stringency, filter criteria, clustering and annotation strategies, the numbers and the corresponding fraction of the genome reported as” enhancers”, were highly variable among the tools (Table 3). Consequently, for the sake of a fair comparison, a comparative analyses at two levels a.1) base pairs a.2) number of enhancers, was performed using the following performance measures. Let A be the number of bases in the predicted enhancers that overlap with the evaluation category of choice, B represents the total number of bases in the predicted enhancers, C denotes the total number of bases in a given evaluation category, n be the number of enhancers overlapped with given evaluation category and N provides the total number of putative enhancers provided at genome-wide level, the mathematical form of the measures can be given as

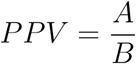

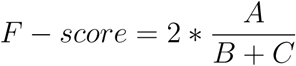

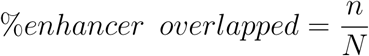

b) Specific signatures of enhancer activity: A comprehensive gold-standard set of experimentally validated enhancers was assembled from different resources corresponding to K562 cell type. The dataset includes 1,621 nonactive enhancer elements, which have shown no activity in CRE-seq assay and 2,392 enhancers, whose activity was confirmed by one of the experimental methods such as VISTA [20] (reporter assay), FANTOM5 [18] (CAGE) and Kwasnieski et al.,[37] (CRE-seq with P-value *<* 0.05). Using this comprehensive set of 1,621 negative (class 0) and 2,392 positive samples (class 1), genome-wide enhancer predictions corresponding to different tools were evaluated using the weighted F1-score (F1” weighted”). To capture the predictive power for positive and negative classes, the weighted F1-score (F1” weighted”) using F1-score of individual classes (F1” 1” and F1” 0”) was computed as shown by following equations using metric module of scikit-learn v0.16.0 python library.

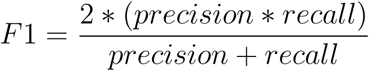

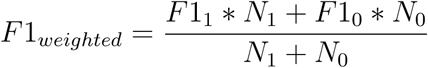

### GEP source and documentation

The computational framework is available at https://github.com/ShaluJhanwar/GEP and the GEP manual has been prepared using readthedocs. Below is a snapshot of GEP online documentation.

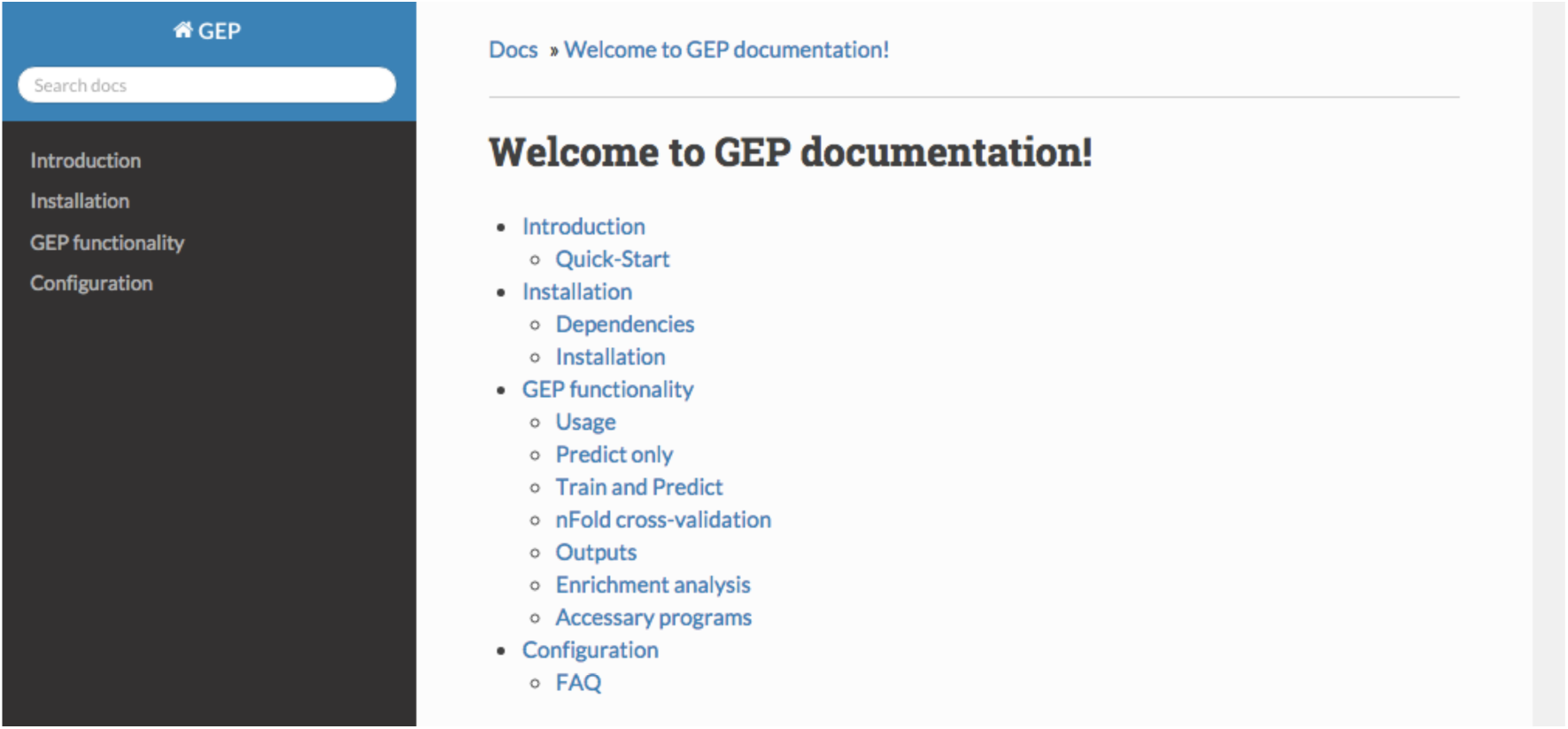

## Author’s contributions

SJ, JD and SO designed the study. SJ implemented the machine-learning models and performed the analysis. SJ, JD and SO wrote the manuscript, contributed ideas and participated in discussions.

## Acknowledgments

We acknowledge support of the Spanish Ministry of Economy and Competitiveness, ‘Centro de Excelencia Severo Ochoa 2013-2017’, SEV-2012-0208. Shalu Jhanwar has been supported by The International PhD scholarship program of La Caixa at CRG. Jose Davilla acknowledges the Centro de Ciencias de la Complejidad (C3), UNAM.

---

**Figure S1:**
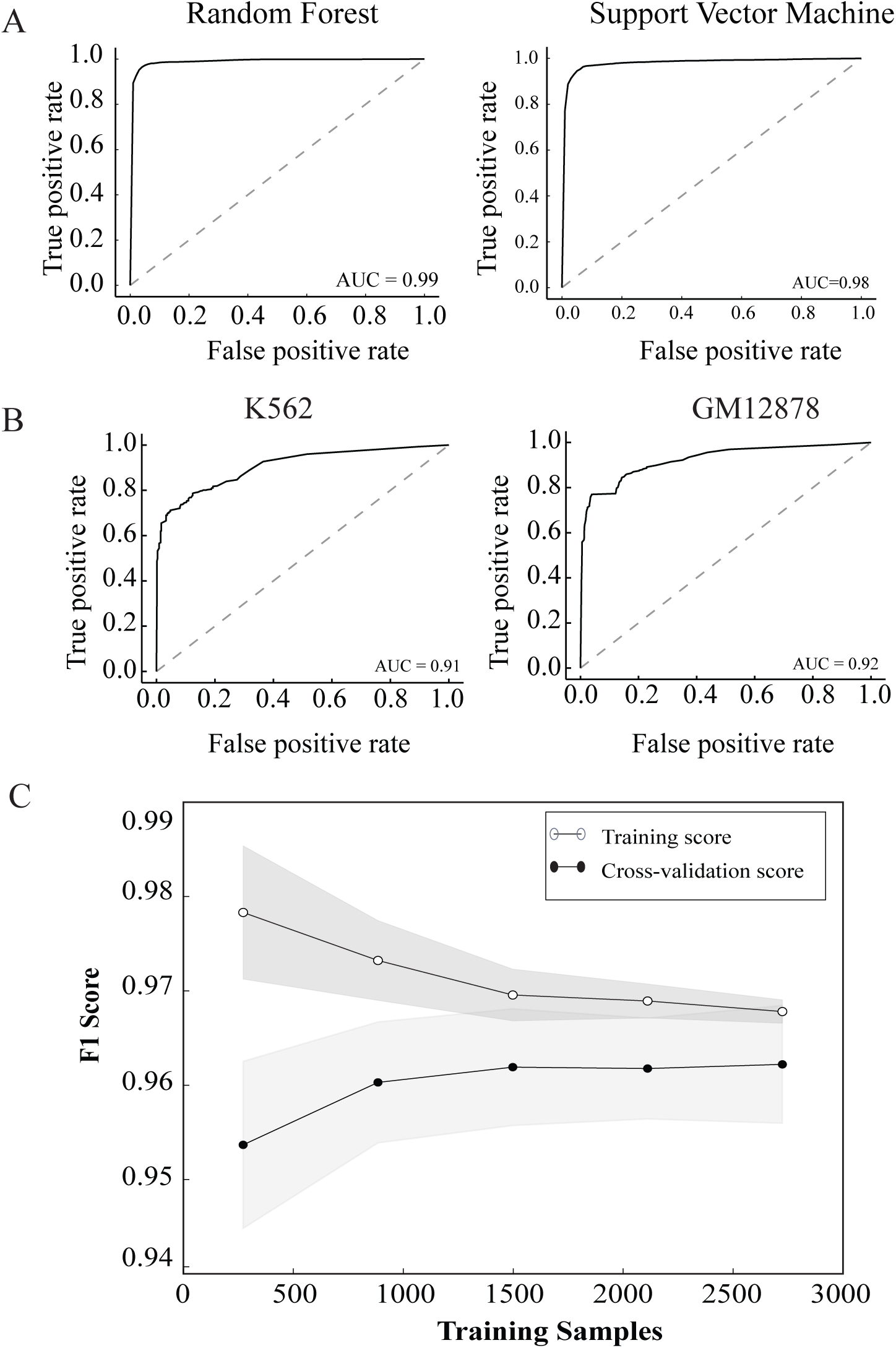
Implementation of classifiers on training and validation datasets. A) ROC representing the mean AUC during 10-fold cross validation of the entire training data B) ROC on validation of FANTOM5 in vivo transcribed enhancer sets using SVM C) Learning curve representing training and testing score in 10 fold stratified cross-validation during training using random forest.

**Figure S2:**
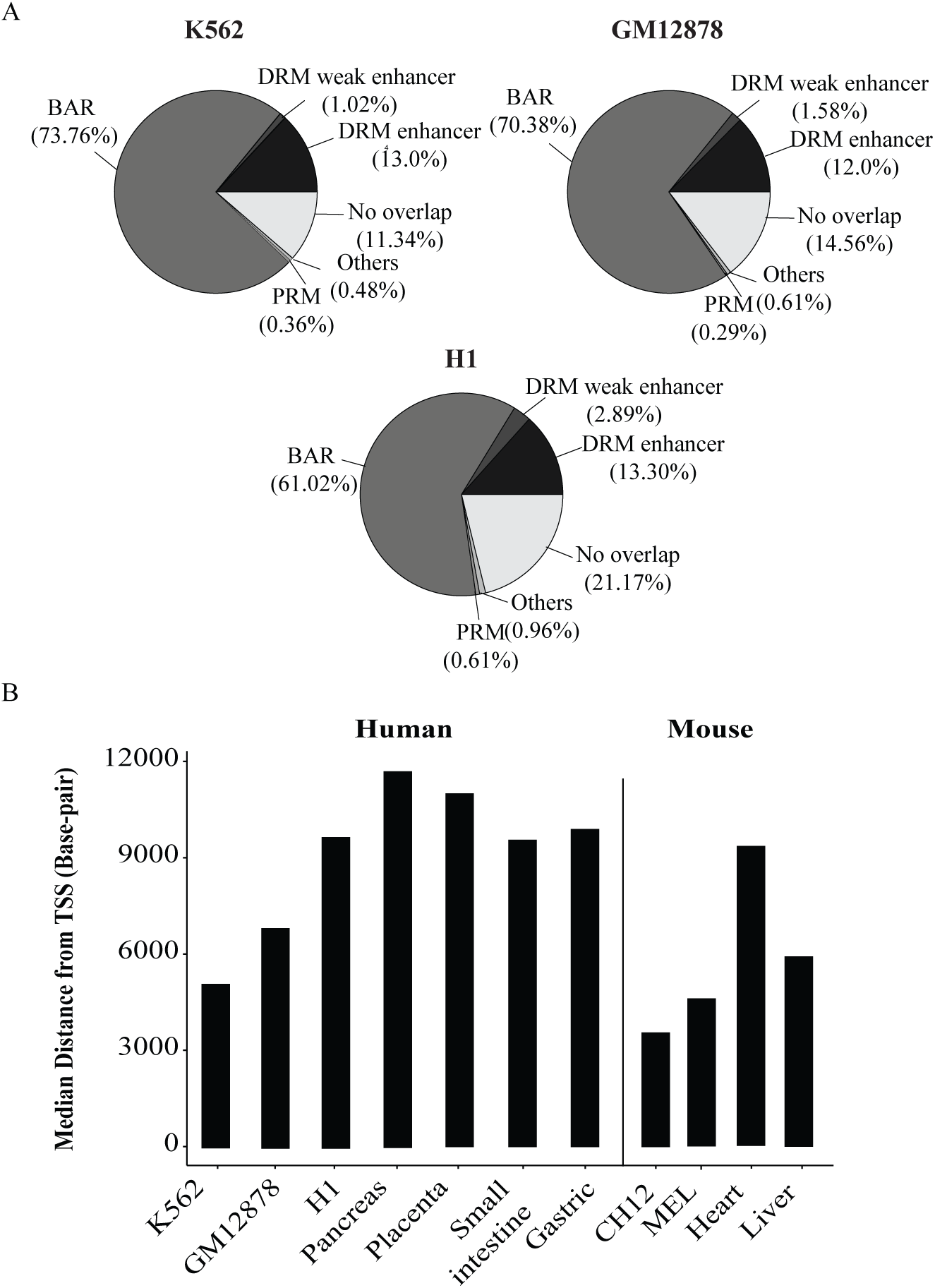
Characterization of genome-wide predicted enhancers using GEP across mammalian cell types and tissues. A) Comparison of predicted enhancers with 6 regulatory modules of Yip et al. [20] predictions, where different modules correspond to distal regulatory module (DRM) containing distal regulatory elements (enhancers, insulators etc), proximal regulatory module (PRM) containing promoters, binding active regions (BAR) with TF binding active regions B) Median distance to TSS (bp) of genome-wide predicted enhancers across different cell-/tissue-types in humans: K562, GM12878, H1, Gastric, Small-Intestine, Pancreas, Placenta and mouse: MEL, CH12.LX, Liver, Heart.

